# An immune-CNS axis activates remote hippocampal stem cells following Spinal Transection Injury

**DOI:** 10.1101/361535

**Authors:** Sascha Dehler, Pak-Kin Lou, Maxim Skabkin, Sabrina Laudenklos, Andreas Neumann, Ana Martin-Villalba

## Abstract

External stimuli such as injury, learning, or stress influence the production of neurons by neural stem cells (NSCs) in the adult mammalian brain. These external stimuli directly impact stem cell activity by influencing areas directly connected or in close proximity to the neurogenic niches of the adult brain. However, very little is known on how distant injuries affect NSC activation state. In this study we demonstrate that a thoracic spinal transection injury activates the distally located hippocampal-NSCs. This activation leads to a transient increase production of neurons that functionally integrate to improve animal’s performance in hippocampal-related memory tasks. We further show that interferon-CD95 signaling is required to promote injury-mediated activation of remote NSCs. Thus, we identify an immune-CNS axis responsible for injury-mediated activation of remotely located NSCs.

## Introduction

The process of generating new neurons in the adult mouse brain is best characterized in the ventricular-subventricular zone (V-SVZ) and the subgranular zone (SGZ) of the dentate gyrus. Neural stem cells (NSCs) within the V-SVZ generate neuronal precursors that migrate along the rostral migratory stream into the olfactory bulbs (OBs) where they disperse radially and generate functional interneurons that fine-tune odor discrimination. NSCs within the SGZ generate neuronal precursors that migrate short distance into the inner granule cell layer of the dentate gyrus where they become functionally integrated into the existing network (Gage, 2000; Taupin & Gage, 2002; Zhao *et al.*, 2008; Ming & Song, 2011; Aimone *et al.*, 2014; Lim & Alvarez-Buylla, 2016). Hippocampal newborn neurons contribute to the formation of certain types of memories such as episodic and spatial memory (Kropff *et al.*, 2015), as well as regulation of mood (Sahay & Hen, 2007) or stress (Snyder *et al.*, 2011; Anacker *et al.*, 2018). Adult neurogenesis is increased by various stimuli like an enriched environment, running and learning via neurotransmitters, hormones or growth factors (Kempermann *et al.*, 1997, 2002; van Praag, Christie, *et al.*, 1999; van Praag, Kempermann, *et al.*, 1999; Nilsson *et al.*, 1999; Shors *et al.*, 2001; van Praag *et al.*, 2005; Leuner *et al.*, 2006; Lledo *et al.*, 2006; Kobilo *et al.*, 2011; Mustroph *et al.*, 2012; Alvarez *et al.*, 2016). In addition, endogenous NSCs can be activated by traumatic brain injury (Arvidsson *et al.*, 2002; Parent *et al.*, 2002; Thored *et al.*, 2006; Hou *et al.*, 2008; Liu *et al.*, 2009).

In this study we show that injury of the spinal cord transiently activates distantly located hippocampal stem cells. Some activated stem cells generate neurons in the hippocampal dentate gyrus that transiently improve performance of injured mice in spatial memory tasks as compared to uninjured controls. Other SGZ-stem cells are activated to migrate away from the dentate gyrus. Notably, we identify the interferon-gamma/CD95 signaling as necessary for activation of NSCs by a remote injury. In summary, our study unveils an immune-CNS interaction leading to injury-mediated activation of hippocampal neurogenesis.

## Material and Methods

### Animals

For the experiments we used the following mouse lines: C57BL/6N, NesCreER^T2^CD95flox [B6.Cg-Tg(Nestin-Cre/Ers1)#GSc Fastm1Cgn] and IFNα-/IFNγ-R-KO [B6.Cg.Ifnar1tm1Agt IfngrltmlAgt / Agt]. Six weeks old NesCreER^T2^ CD95flox (Cre^+^) and respective controls (Cre^-^) were intraperitoneally (i.p.) injected with 1mg Tamoxifen (Sigma) twice a day for 5 consecutive days before operating. At the age of 12 weeks the respective group of mice received a sham or spinal transection injury as previously described (Letellier *et al.*, 2010). For short term labelling of NSCs, mice received i.p. BrdU (Sigma;300mg/kgbw) injections at 1h, 24h and 48h post injury or a single shot injection 89 days post injury (Figure 1A, Figure 4A and Figure 4E), followed by a chase time of 1 day, 2 weeks or 4 weeks, respectively. For the long term label retaining experiment (Figure 2A), 8 weeks old mice received a daily single shot injection of BrdU (50mg/kgbw) for a total duration of three weeks followed by a chase time of 16 weeks after the last BrdU injection. For the isolation of primary neural stem cells, 8 weeks old C57BL/6N mice were used. All animals were housed in the animal facilities of the German Cancer Research Center (DKFZ) at a 12 hrs. dark/light cycle with free access to food and water. For the injury and behavioral experiments, exclusively age-matched female mice were used. All animal experiments were performed in accordance with the institutional guidelines of the DKFZ and were approved by the “Regierungspräsidium Karlsruhe”, Germany.

**Figure 1:**
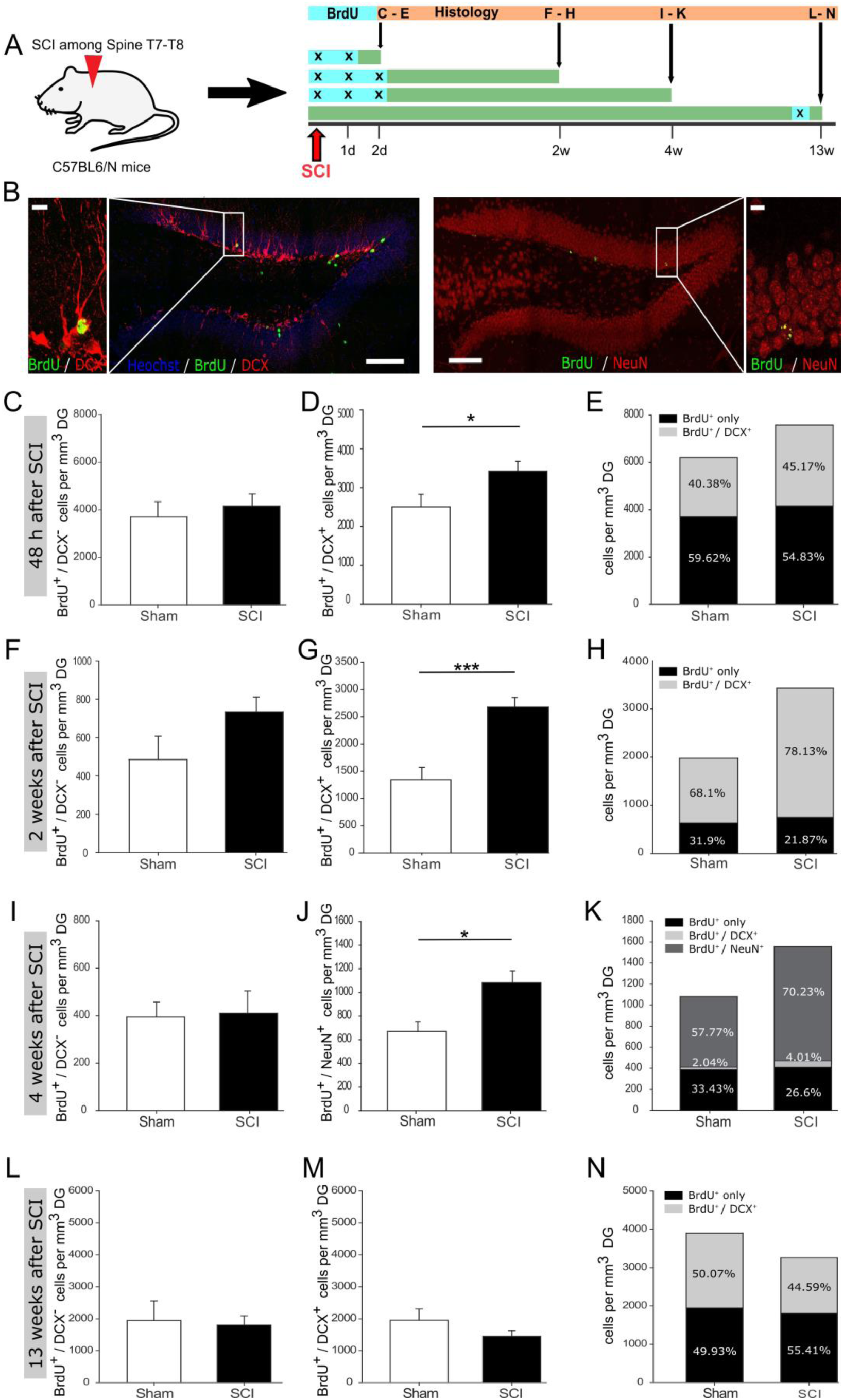
Increased hippocampal neurogenesis upon distant spinal cord injury. (A) Schematic illustration of experimental timeline performed with C57BL/6N mice. (B) BrdU incorporation within the dentate gyrus of adult mice. Scale bar is 100 μm or 10 μm, respectively. (C) Quantification of BrdU^+^/DCX^-^ cells 48 h post injury, mean values (± SEM) from sham (3698 ± 561 cells/mm^3^ DG) vs. SCI (4156 ± 434 cells/mm^3^ DG) mice, group size n_sham_ =6 vs. n_SCI_ =6. (D) Quantification of BrdU^+^/DCX^+^ cells 48 h post injury, mean values (± SEM) from sham (2505 ± 323 cells/mm^3^ DG) vs. SCI (3422 ± 249 cells/mm^3^ DG) mice, group size n_sham_ =6 vs. n_SCI_ =6, ^∗^p < 0.05 (Student’s t-test). (E) Percentage distribution of BrdU^+^/DCX^+^ cells in sham (40.38%) and SCI (45.17%) mice, 48 h following injury. (F) Quantification of BrdU^+^/DCX^-^ cells two weeks post injury, mean values (+SEM) from sham (485 ± 109 cells/mm^3^ DG) vs. SCI (734 ± 70 cells/mm^3^ DG) mice, group size n_sham_ =5 vs. n_SCI_ =6. (G) Quantification of BrdU^+^/DCX^+^ cells two weeks post injury, mean values (± SEM) from sham (1345 ± 224 cells/mm^3^ DG) vs. SCI (2677 ± 175 cells/mm^3^ DG) mice, group size n_sham_ =6 vs. n_SCI_ =6, ^∗∗∗^p < 0.001 (Student’s t-test). (H) Percentage distribution of BrdU^+^/DCX^+^ cells in sham (68.1%) and SCI (78.13%) mice, two weeks following injury. (I) Quantification of BrdU^+^/DCX^-^ cells four weeks post injury, mean values (± SEM) from sham (394 ± 57 cells/mm^3^ DG) vs. SCI (410 ± 86 cells/mm^3^ DG) mice, group size n_sham_ =5 vs. n_SCI_ =6. (J) Quantification of BrdU^+^/NeuN^+^ cells four weeks post injury, mean values (± SEM) from sham (669 ± 83 cells/mm^3^ DG) vs. SCI (1082 ± 99 cells/mm^3^ DG) mice, group size n_sham_ =6 vs. n_SCI_ =6, ^∗^p < 0.05 (Student’s t-test). (K) Percentage distribution of BrdU^+^/NeuN^+^ cells in sham (57.77%) and SCI (70.23%) mice and BrdU^+^/DCX^+^ cells in sham (2.04%) and SCI (4.01%) mice, four weeks following injury. (L) Quantification of BrdU^+^/DCX^-^ cells 13 weeks post injury, mean values (± SEM) from sham (1947 ± 558 cells/mm^3^ DG) vs. SCI (1805 ± 270 cells/mm^3^ DG) mice, group size n_sham_ =6 vs. n_SCI_ =8. (M) Quantification of BrdU^+^/DCX^+^ cells 13 weeks post injury, mean values (± SEM) from sham (2005 ± 382 cells/mm^3^ DG) vs. SCI (1452 ± 159 cells/mm^3^ DG) mice, group size n_sham_ =5 vs. n_SCI_ =8. (N) Percentage distribution of BrdU^+^/DCX^+^ cells in sham (50.07%) and SCI (44.59%) mice, 13 weeks following injury.

**Figure 2:**
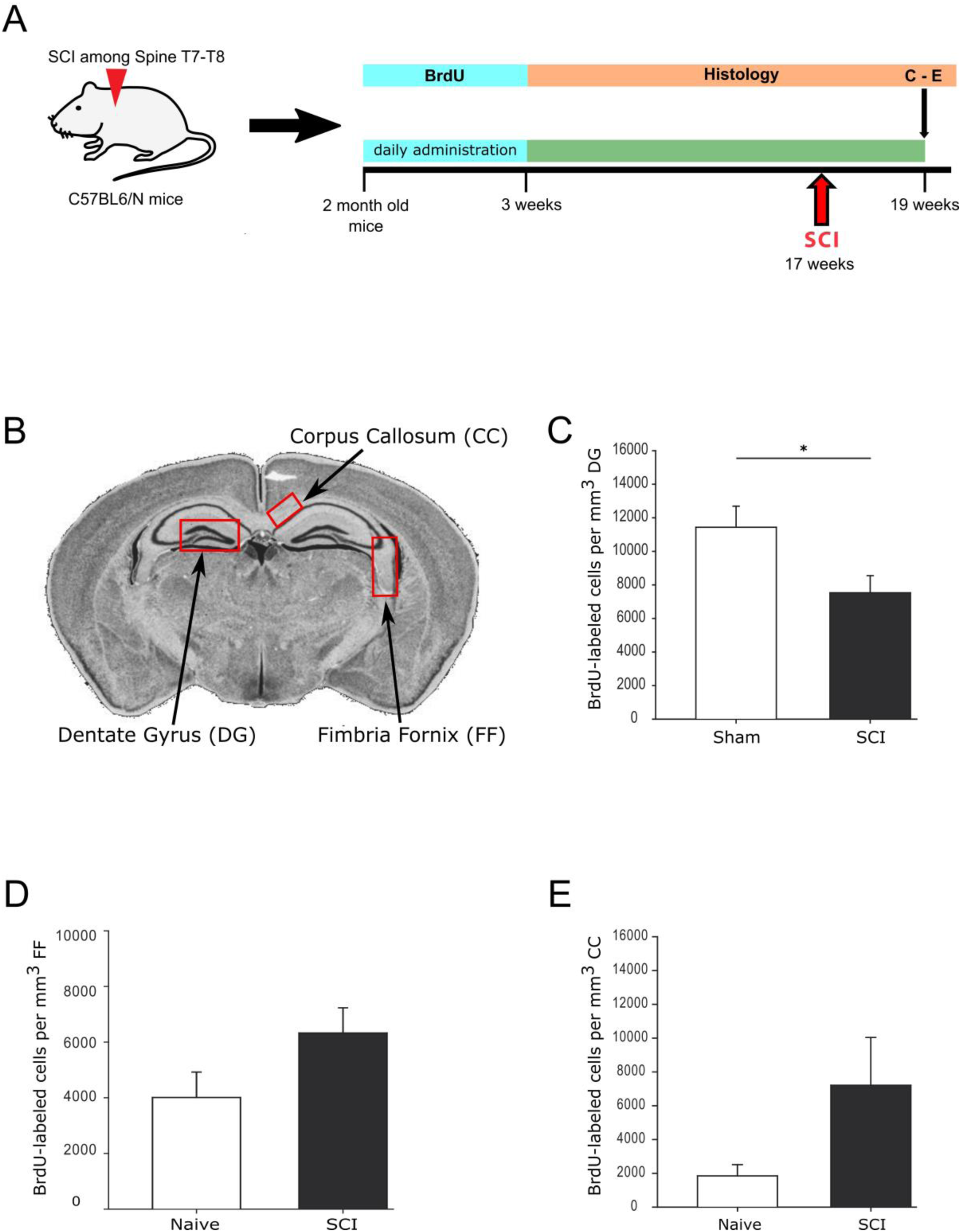
Dormant NSCs are activated following spinal transection injury. (A) Schematic illustration of the experimental timeline for labeling dormant NSCs in the DG of adult C57BL/6N mice. (B) Representative coronal section of the adult mouse brain with designated regions for the quantification of BrdU^+^ labeled cells. (C) Quantification of BrdU^+^ labeled cells in the DG, mean values (± SEM) from sham (11437 ± 1255 cells/ mm^3^ DG) vs. SCI (7532 ± 1017 cells /mm^3^ DG) mice, group size n_sham_ =5 vs. n_SCI_=11, ^∗^p < 0.05 (Student’s t-test). (D) Quantification of BrdU^+^ labeled cells in the FF, mean values (± SEM) from naive (4010 ± 913 cells/ mm^3^ FF) vs. SCI (6326 ± 906 cells /mm^3^ FF) mice, group size n_naive_ =4 vs. n_SCI_ =6. (E) Quantification of BrdU^+^ labeled cells in the CC, mean values (± SEM) from naive (1848 ± 665 cells/ mm^3^ CC) vs. SCI (7210 ± 2829 cells /mm^3^ CC) mice, group size n_naive_ =4 vs. n_SCI_ = 6

### Spinal Cord Injury

Female, age-matched animals were subjected to laminectomy at spine T7-T8 followed by a 80% transaction of the spinal cord injury by cutting the spinal cord with iridectomy scissors, as described in (Demjen *et al.*, 2004; Stieltjes *et al.*, 2006; Letellier *et al.*, 2010). Sham mice were subjected only to laminectomy. Naïve mice did not face any surgical procedure.

### Handling of the Animals

Mice were habituated to the handling experimenter before starting with behavioural experiments. To this end, mice were handled for 5-10 minutes twice a day. Handling was performed for at least 5 days until the animals showed no anxiety-related behaviour when meeting the experimenter.

### Spontaneous alternation in the T-maze

Spatial working memory performance was assessed on an elevated wooden T-Maze as described in (Corsini *et al.*, 2009). Each animal had 4 sessions on the T-Maze (1 session/day; 4 trials/session). One trial consisted of a choice and a sample run. During the choice run one of the two target arms was blocked by a barrier according to a pseudorandom sequence, with equal numbers of left and right turns per session and with no more than two consecutive turns in the same direction. The mice were allowed to explore the accessible arm. Before the sample run (intertrial interval of ~10 sec) the barrier was removed enabling accessibility to both arms. On the sample run the mouse was replaced back into the start arm facing the experimenter. The mouse was allowed to choose one of the two target arms. The trial was classified as success if the animal chose the previously blocked arm. For analysis all trials were combined and the success rate (%) was quantified ((# successful trials/# trials)^∗^100).

### Immunohistochemistry

Animals were sacrificed by using an overdose of Ketamin (120mg/kg) / Xylazine (20mg/kg) and were subsequently transcardially perfused with 20ml 1xHBSS (Gibco) and 10ml of 4% paraformaldehyde (Carl Roth). The brains were dissected and postfixed in 4% paraformaldehyde overnight at 4 °C. A Leica VT1200 Vibratome was used to cut the tissue in 50μm thick coronal sections. From each mouse six Brain sections every 300μm along the coronal axis were used for quantification. First, the brain sections were washed 3× 15 min at room temperature in TBS, followed by a 1hrs blocking step in TBS^++^ (TBS with 0.3% horse serum (Millipore) and 0.25% Triton-X100 (Sigma)) at room temperature. Tissue was transferred to 0.5ml Safe Lock Reaction-Tubes containing 200μl TBS^++^ including primary antibodies. Samples were incubated for 24-48 hours at 4°C. After incubating with primary antibody, tissue samples were washed 3× 15min in TBS at room temperature, followed by a 30 min blocking step in TBS^++^ at room temperature. Brain sections were transferred to 0.5ml Safe Lock Reaction-Tubes containing 200μl TBS^++^ including secondary antibodies. Samples were incubated in the dark, for 2 hrs at room temperature. Finally the brain slices were washed 4× 10 min in TBS at room temperature, before they were further floated in 0.1M PB-Buffer and mounted on glass slides with Fluoromount G (eBioscience). The following antibodies were used: rat anti-BrdU (Abcam, 1/150), goat anti-DCX (Santa Cruz, 1/200) and mouse anti-NeuN (Merck Millipore, 1/800). Nuclei were counterstained with Hoechst 33342 (Biotrend, 1/4000).

### Microscopy and Cell Quantification

All images were acquired with a Leica TCS SP5 AOBS confocal microscope (Leica) equipped with a UV diode 405nm laser, an argon multiline (458-514nm) laser, a helium-neon 561nm laser and a helium-neon 633nm laser. Images were acquired as multichannel confocal stacks (Z-plane distance 2μm) in 8-bit format by using a 20× (HCX PL FLUOTAR L NA0.40) oil immersion objective. Images were processed and analyzed in ImageJ (NIH). For representative images, the maximum intensity of a variable number of Z-planes was stacked, to generate the final Z-projections. Representative images were adjusted for brightness and contrast, applied to the entire image, cropped, transformed to RGB color format and assembled into figures with Inkscape. For cell quantification the entire volume of the DG was calculated by multiplying the entire area of the DG (middle plane of the total Z-stack) with the entire Z-stack size. The different cell populations were identified and counted (LOCI and Cell-Counter pug-in for ImageJ) based on their antibody labeling profile. Cell counts were either represented as Cells/mm^3^DG or as Cells/DG.

### *In vitro* culturing and treatment of NSCs with INFγ

The lateral SVZ was microdissected as a whole mount as previously described (Mirzadeh *et al.*, 2010). Tissue of one mouse was digested with trypsin and DNase according to the Neural Tissue Dissociation Kit (Miltenyi Biotec) in a Gentle MACS Dissociator (Miltenyi Biotec). Cells were cultured and expanded for 8-12 days in Neurobasal medium (Gibco) supplemented with B27 (Gibco), Heparine (Sigma), Glutamine (Gibco), Pen/Strep (Gibco), EGF (PromoKine) and FGF (PeloBiotech) as used in (Walker & Kempermann, 2014). For stimulation with INFγ (Millipore), 4×10^5^ cells were seeded. The next day, cells were treated with 50ng INFγ / ml media for duration of 14 hours.

### Flow Cytometric Analysis

The cells were harvested and were treated with Accutase (Sigma) for 5 min at 37 °C, followed by filtering the cells with a 40μm cell strainer to get a single cell suspension. Afterwards the cells were washed twice with FACS media (PBS/10%FCS) and were re-suspend in 200μl FACS media. Cells were stained for 30 min at room temperature by using the Jo2 CD95::PECy7 antibody (BD Pharming/ 1/100). Afterwards the cells were washed three times with FACS media and were finally re-suspend in 200μl FACS media.

## Results

### Enhanced Hippocampal neurogenesis following spinal cord injury

To assess whether a remote CNS injury would activate NSCs in the SGZ, we injured the spinal cord at thoracic level T7-T8. In order to detect the reaction of SGZ-NSCs and their neurogenic progeny, we labeled these cells with 5-Bromo-2-deoxyuridine (BrdU; once daily) at the time of injury and in the following 24h, 48h or after 89 days and examined them at 2d, 2 weeks, 4 weeks and 13 weeks following injury (Figure 1A). Brains were stained for BrdU, to follow actively dividing NSCs (BrdU), quiescent NSCs and neuroblasts (Figure 1B). Already 48 hours after injury, we observed a significant increase in new-born neuroblasts (BrdU^+^/DCX^+^) and an increased number of BrdU^+^/DCX^-^ cells (Figure 1C-E). 2 weeks following injury, the number of BrdU^+^/DCX^-^ cells and neuroblasts was significantly higher when compared to sham-injured controls (Figure 1F-H). We further assessed the maturation of BrdU-labelled cells to neurons (BrdU^+^/NeuN^+^) at 4 weeks after the injury. Significantly more newborn neurons were identified in the dentate gyrus of the injured mice, whereas the number of BrdU^+^/DCX^-^ cells was comparable in injured and sham controls (Figure 1I-K). Notably, at 13 weeks following injury, the number of cycling BrdU^+^/DCX^-^ cells and newborn neuroblasts was set back to basal levels, exhibiting similar numbers to that of its sham operated counterparts (Figure 1L-N). Together, our data showed that distant spinal cord injury stimulates a fast but transient activation of NSCs in the remote SGZ to generate neurons.

### Spinal Cord Injury induced migration out of the dentate gyrus of dormant SGZ-NSCs

Injury has been shown to activate a pool of highly dormant cells in the hematopoietic system (Wilson *et al.*, 2008; Essers *et al.*, 2009; Essers & Trumpp, 2010). To test if this is also the case for SGZ-NSCs, we used a three weeks BrdU-labelling protocol starting at the age of 8-weeks and allowed a chase time of 16 weeks after the last BrdU injection. Mice were subjected to spinal cord injury at 14 weeks chase time or left uninjured and sacrificed two weeks later to follow the reaction to injury of the highly dormant NSCs (Figure 2A). Notably, the number of BrdU^+^ cells in the DG was significantly reduced in injured mice as compared to sham controls (Figure 2B-C). The default program for SGZ-NSCs is to become active and produce neuroblasts that further mature to granule cell neurons within the dentate gyrus. Yet, it is known that upon injury some NSCs in close vicinity start migrating out of the neurogenic niche (Nakatomi *et al.*, 2002; Grande *et al.*, 2013). Therefore we assessed a potential migration of BrdU-labelled cells to the neighboring regions of the fimbria-fornix (FF) and corpus callosum (CC) (Figure 2B). The quantification of BrdU^+^ cells in the FF and the CC showed a trend to an increased number of BrdU-labelled cells in these nearby regions in injured mice as compared to naïve counterparts (Figure 2D-E). In summary our data suggest that spinal cord injury activates alternative migratory pathways in a pool of highly dormant, long-term label-retaining SGZ-NSCs.

### Spinal Cord Injury leads to better Working Memory

Together we see that the injury activates both, normal homeostatic neurogenesis and decreases the pool of highly dormant stem cells potentially by activating their migration out of the dentate gyrus. Therefore, we next tested the function of the injury-induced surplus of newborn neurons within the hippocampus of injured mice, homeostatic neurogenesis. Adult hippocampal neurogenesis has been shown to positively impact short- and long-term spatial working memory, navigation learning, pattern discrimination as well as trace and contextual fear conditioning (Corsini *et al.*, 2009 Deng *et al.*, 2010; Aimone *et al.*, 2011), but also to counteract depression– and stress-induced behavioral responses (Sahay & Hen, 2007; Snyder *et al.*, 2011). To test the function of injury-induced newborn neurons in the dentate gyrus, we tested the performance of injured and naïve mice in a hippocampal-dependent task, the spontaneous alternation on an elevated T-Maze, used as readout of short term spatial working memory (Figure 3A). Mice were tested at two, four and eight weeks following spinal cord injury. At two weeks post-injury naïve, sham and spinal cord injured mice showed a similar success rate of the spontaneous alternation (Figure 3B). Importantly, at four weeks following injury, the success rate of injured mice was significantly higher than the rate of Sham controls (Figure 3C). Notably, the improved performance of injured mice on the T-Maze disappeared at eight weeks post-injury (Figure 3D). Taken together, our data demonstrated that newly generated neurons integrate into the existing hippocampal network and positively influence the performance of injured mice in a hippocampal-dependent spatial memory task. However, as the observed activation of neurogenesis, the functional improvement is also transient. Interestingly, we previously observed a transient increase in neurogenesis following exercise that improved performance on the T-Maze in an equally transient mode (Corsini *et al.*, 2009). Taken together, our data suggest that the newborn functionally immature neurons impact short term memory as long as they are young and plastic. However, this effect disappears as they become similar to their older counterparts (Kropff *et al.*, 2015).

**Figure 3:**
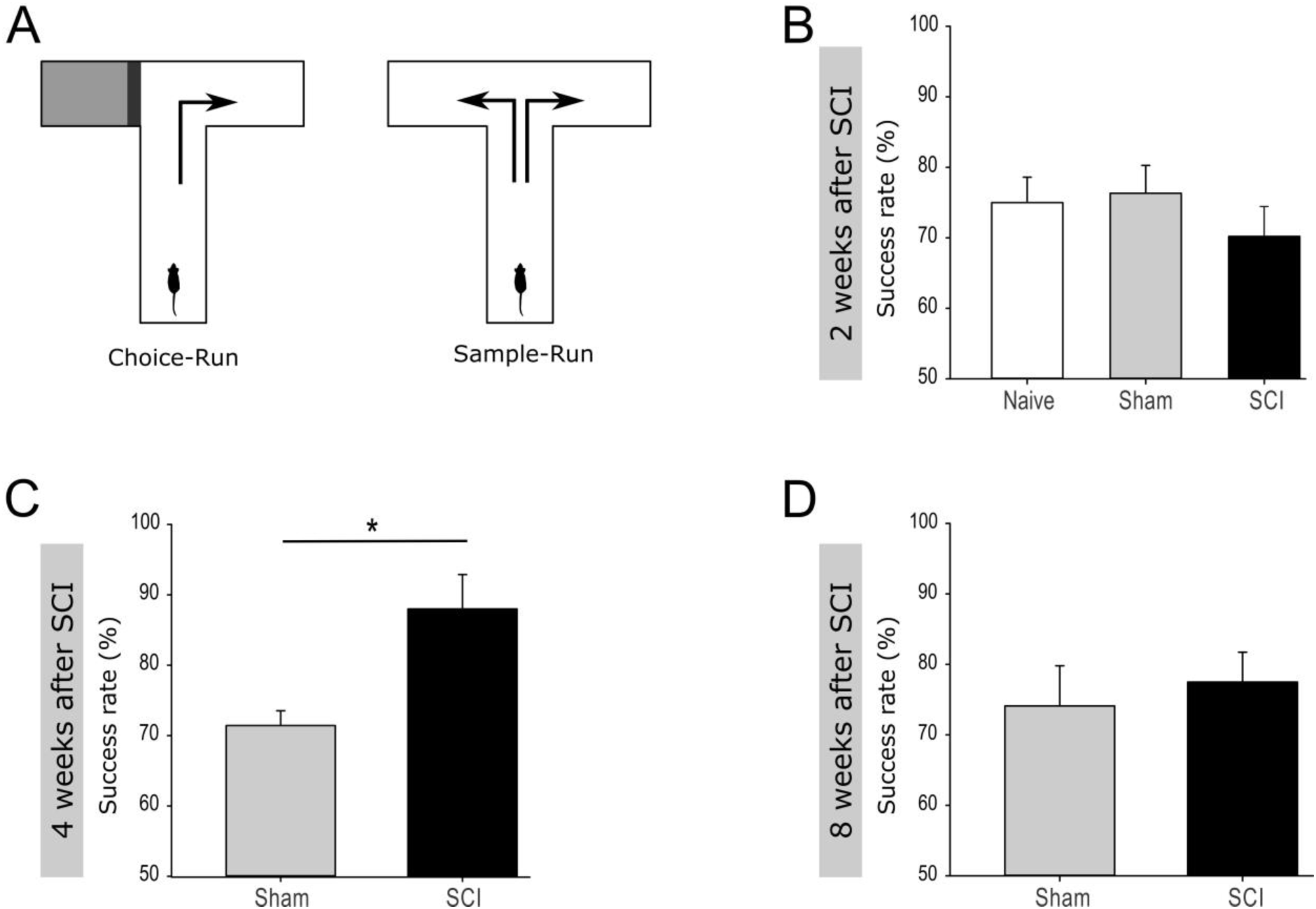
Improved performance in a Working Memory task following spinal cord injury. (A) Experimental setup for the spontaneous alternation in the T-Maze test. (B) Mean success rate (± SEM) of naïve (75% ± 3.29) vs. sham (76.34% ± 3.80) vs. SCI (70.19% ± 4.09) mice, two weeks post injury, group size n_naive_ =6 vs. n_sham_ =14 vs. n_SCI_=13. (C) Mean success rate (± SEM) of sham (71.43% ± 5.15) vs. SCI (88% ± 4.38) mice, four weeks post injury, group size, n_sham_ =7 vs. n_SCI_ =5, ^∗^p < 0.05 (Mann-Whitney test). (D) Mean success rate (± SEM) of sham (74.11% ± 5.27) vs. SCI (77.5% ± 3.79) mice, eight weeks post injury, group size, n_sham_ =7 vs. n_SCI_ =5.

### Loss of IFNα-/IFFNγ-R and CD95 inhibits neural stem cell activation upon spinal transection injury

Acute tissue injury activates an immediate inflammatory response that is able to rapidly affect distant locations. Notably, we previously identified interferons as an activator of NSCs in the V-SVZ following a global ischemic insult that induces damage in the nearby located striatum (Llorens-Bobadilla *et al.*, 2015). The requirement of IFNγ signalling for SCI-mediated activation of SGZ-NSCs was further tested using mice deficient in IFNα-/IFNγ-receptor as compared to wt counterparts (Figure 4A, Figure 1F-H). Excitingly, two weeks following injury, IFNα-/IFNγ-receptor deficient mice did neither show a significant increase in the population of neuroblasts (BrdU^+^/DCX^+^), nor in the population of BrdU^+^/DCX^-^ cells (Figure 4B-D). These observations indicated that spinal transection injury does not activate SGZ-NSCs lacking a functional IFNα/IFNγ-signaling-pathway. We next investigated the putative signalling pathways involved in local SCI-mediated activation of SGZ-NSCs. Interferons have been reported to increase the expression of CD95-ligand and CD95 (Chow *et al.*, 2000; Kirchhoff *et al.*, 2002; Boselli *et al.*, 2007). In a previous study we demonstrated that the TNF-R family member, CD95, is required for the activation of SGZ-NSCs following global ischemia (Corsini *et al.*, 2009). To test the regulation of CD95 upon IFNγ treatment in NSCs, we isolated NSCs from the V-SVZ of 8 weeks old C57BL/6N mice, cultured them in vitro for short time and exposed them for 14 hours to IFNγ. Thereafter expression of CD95 was analysed by Flow Cytometry. IFNγ significantly increased the expression of CD95 in NSCs as compared to untreated NSCs (Figure 4H and Supplementary Figure S1). To assess CD95’s involvement in SCI-induced neurogenesis we used the NesCreER^T2^CD95^f/f^ mouse line. This mouse line enables an acute deletion of CD95 in the adult neural stem cell compartment (Corsini *et al.*, 2009). CD95NesCreER^T2+^ (Cre) and CD95NesCreER^T2-^ (Cre^-^) mice received tamoxifen injections at 6 weeks of age. Their spinal cord was injured at the age of 12 weeks. Dividing cells were labelled by BrdU at the time of injury and 24 hours post injury. The SGZ was further processed for staining of BrdU and DCX 48 hours after the surgery (Figure 4E). CD95-deficient NSCs exhibit an impaired injury-induced activation, as fewer BrdU^+^/DCX^-^ cells and newborn neuroblasts (BrdU^+^/DCX^+^) could be detected in the SGZ of Cre^+^ mice as compared to their injured Cre^-^ counterparts (Figure 4F-G). Thus, CD95 is locally involved in activation of SGZ-NSCs by a remote injury. Next, we set out to test if the injury-induced improvement of the spatial working-memory is due to the increased activation of NSCs. Indeed, injured and sham operated IFNα-/IFNγ-receptor deficient mice showed a similar success rate in the spontaneous alternation in the elevated T-Maze (Figure 4I-K). Thus, interferon-related increased of homeostatic neurogenesis mediates the functional improvement in short-term working memory exhibited by spinal injured animals. Altogether, our results indicate that injury-induced IFN signaling triggers CD95 activation of SGZ-NSCs, thereby leading to a transient expansion of the pool of newborn neurons resulting in an improved working memory.

**Figure 4:**
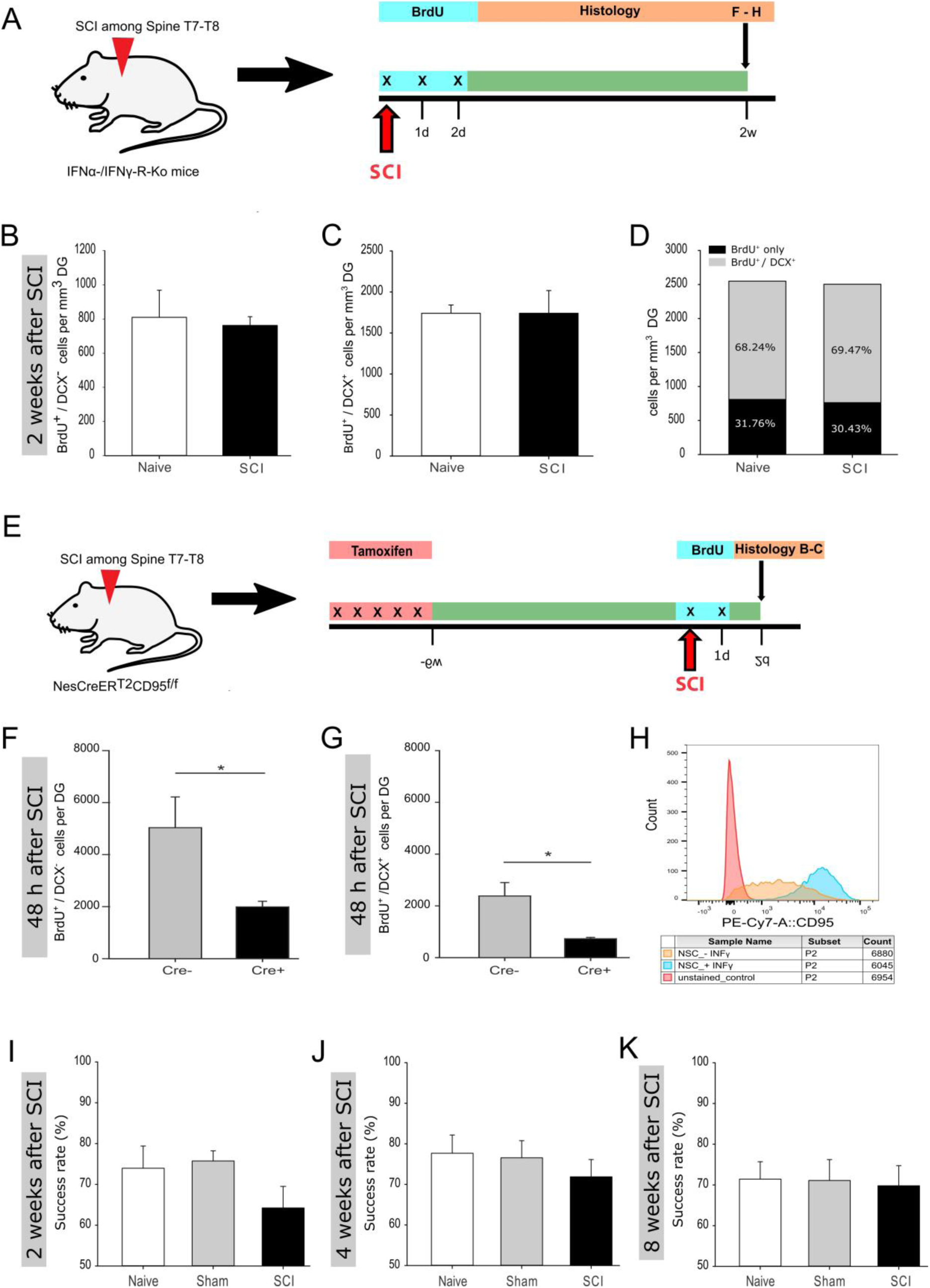
Reduced activation of adult hippocampal neurogenesis in IFNα-IFNγ-R and CD95-Ko upon spinal cord injury. (A) Illustration of the experimental timeline performed with IFNα-/IFNγ-R-Ko mice. (B) Quantification of BrdU^+^/DCX^-^ cells two weeks post injury, mean values (± SEM) from naïve (809 ± 137 cells/mm^3^ DG) vs. SCI (762 ± 46 cells/mm^3^ DG) mice, group size n_naive_ =4 vs. n_SCI_ =5. (C) Quantification of BrdU^+^/DCX^+^ cells two weeks post injury, mean values (± SEM) from naïve (1740 ± 88 cells/mm^3^ DG) vs. SCI (1741 ± 247 cells/mm^3^ DG) mice, group size n_naive_ =4 vs. n_SCI_ =5. (D) Percentage distribution of BrdU^+^/DCX^+^ cells in naïve (68.24%) and SCI (69.47%) mice, two weeks following injury. (E) Illustration of the experimental timeline performed with NesCreER^T2^CD95^f/f^ mice. (F) Quantification of BrdU^+^/DCX^-^ cells 48 h post injury, mean values (± SEM) from injured Cre^-^ (5034 ± 2899 cells/DG) vs. injured Cre^+^ (1989 ± 530 cells/DG) mice, group size n_Cre_- =6 vs. n_Cre_+ =6, ^∗^p < 0.05 (Student’s t-test). (G) Quantification of BrdU^+^/DCX^+^ cells 48h post injury, mean values (± SEM) from injured Cre^-^ (2381 ± 1273 cells/DG) vs. injured Cre^+^ (728 ± 126 cells/DG) mice, group size n_Cre_- =6 vs. n_Cre_+ =6, ^∗^p < 0.05 (Student’s t-test). (H) Relative CD95 expression in unstained control, INFγ-untreated and -treated cells are illustrated in a single parameter histogram. (I) Mean success rate (± SEM) of naïve (73.96% ± 4.98) vs. sham (75.75% ± 2.33) vs. SCI (64.22% ±4.62) mice, two weeks post injury, group size n_naive_ =6 vs. n_sham_ =8 vs. n_SCI_ =8. (J) Mean success rate (± SEM) of naïve (77.68% ± 4.49) vs. sham (76.56% ± 3.95) vs. SCI (71.88 ± 3.98) mice, four weeks post injury, group size n_naive_ =7 vs. n_sham_ =8 vs. n_SCI_ =8. (K) Mean success rate (± SEM) of naïve (71.43% ± 4.28) vs. sham (71.09% ± 4.81) vs. SCI (69.79 ± 3.91) mice, eight weeks post injury, group size n_naive_ =7 vs. n_sham_ =8 vs. n_SCI_ =6.

## Discussion

Here, we examine how a remote injury to the CNS influences distally located SGZ-NSCs, short and long term post-injury. Our data clearly show an acute and transiently increased activation of adult SGZ-NSCs to produce neurons following a remote injury and suggest that a fraction of highly dormant stem cells are activated to migrate out of the neurogenic niche. Notably, we show that the newly generated neurons functionally integrate into the existing network, as demonstrated in an elevated spatial navigation performance. However, this activation of neurogenesis fades away with time. Accordingly, two studies investigated the effects of spinal cord injury to the neurogenic niches in adult *Sprague*-*Dawley* rats and detected a decreased level of adult v-SVZ and SGZ neurogenesis 60 days post spinal cord injury (Felix *et al.*, 2012; Jure *et al.*, 2017). Besides, studies of hematopoietic stem cell (HSC) activation by inflammatory signals, show that an acute exposure activates the quiescent population of HSCs, whereas chronic exposure negatively impact HSC activation (Essers *et al.*, 2009).

As already hypothesized by Felix and colleagues (Felix *et al.*, 2012) and in line with Essers et al. (Essers *et al.*, 2009) we show that inflammatory signatures, released in an acute phase post spinal cord injury, play a major role in transmitting the injury signal towards the hippocampus to activate adult neurogenesis. By using single cell transcriptomics we have shown before that interferon-gamma is an important factor in activating v-SVZ neurogenesis in a model of global ischemia (Llorens-Bobadilla *et al.*, 2015). In the current study, we identify interferons as the main factor that transmits the injury signal from the spinal cord towards the hippocampus, where through activation of CD95 stem cells exit the quiescent state to differentiate into neurons. The observed transition from a quiescent to an active state, triggered by a distant injury site, in effects seems to be similar to the transition from G0 to an elevated G_alert_ state in muscle satellite cells (Rodgers *et al.*, 2014). Interestingly, this alert state is triggered in distant stem cells in contralateral muscles, and is also observed in other tissue stem cells such as hematopoietic stem cells (Rodgers *et al.*, 2014). Stem cells in an alert state are primed for cell cycle entry to react in a much faster and efficient way to incoming injuries of different nature. Here we show that a remote CNS injury triggers different responses in actively dividing and dormant NSCs. While actively dividing NSCs are engaged in homeostasis, the fraction of dormant cells decreases, presumably to take alternative migratory pathways to injury-associated areas. However, future studies are needed to follow up the fate of these highly dormant stem cells.

What could be the role of an increased production of granule cell neurons in the hippocampus? Certainly, spinal cord injury represents a very stressful state for the whole organism. It has been shown that adult hippocampal neurogenesis is on the one hand strongly influenced by chronic and acute stress (Conrad *et al.*, 1999; Kirby *et al.*, 2013; LaDage, 2015), on the other hand increased neurogenesis ameliorates stress (Snyder *et al.*, 2011; Anacker *et al.*, 2018). Thus, we speculate that the elevated levels of neurogenesis upon spinal cord injury might buffer injury-induced stress and enable an improved behavioral adaptation to the post injury situation.

In summary, our data show that an acute injury to the spinal cord activates hippocampal neurogenesis, resulting in a transiently increased production of newborn neurons that are functional, as shown by the improved performance in spatial memory tasks of injured mice. Furthermore, we identified interferons as a major factor involved in activation via CD95 of distant stem cells.

## Acknowledgements

We thank C. Pitzer and the members of the Interdisciplinary Neurobehavioral Core (INBC); M. Essers for Ifngr1^-/-^ and Ifnar^-/-^ mice; S. Limpert, K. Volk and M. Richter for technical assistance; the Light Microscopy Core Facility; the DKFZ Flow Cytometry Core Facility; and the members of the Martin-Villalba laboratory for critically reading the manuscript.

---

**supplemtentary Figure S1, related to Figure 4:**
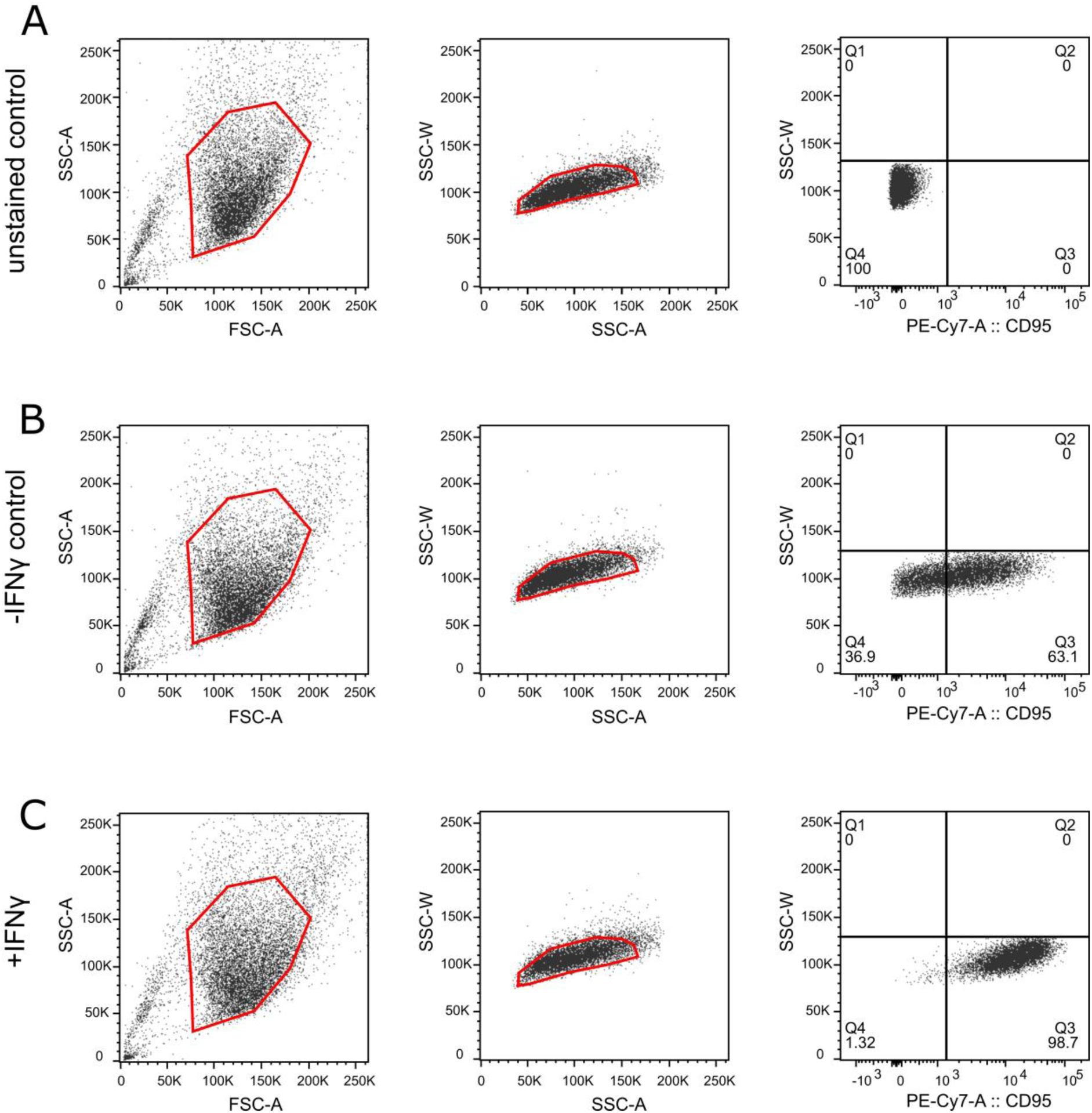
Strategy to determine relative CD95 expression in cultured NSCs by using Flow Cytometry. First gate uses FSC/SSC gating to exclude cellular debris; second gate excludes cell aggregates and third shows relative CD95 expression in unstained control cells (A), stained IFNγ-untreated cells (B) and stained IFNγ-treated cells (C).

## Footnotes

**Author Contributions** S.D. performed experiments, analyzed and interpreted data, and prepared the manuscript. P.L.; M.S.; S.L and A.N. performed experiments and analyzed data. A.M.-V. took over the conceptual design and the study coordination, interpreted data, and prepared the manuscript.

